# Harmonizing the stimulation dose of focal tDCS across target sites

**DOI:** 10.64898/2026.01.09.698549

**Authors:** Axel Thielscher, Dayana Hayek, Oula Puonti, Ulrike Grittner, Felix Blankenburg, Rico Fischer, Gesa Hartwigsen, Shu-Chen Li, Marcus Meinzer, Michael A. Nitsche, Dagmar Timmann, Agnes Flöel, Daria Antonenko

**Affiliations:** Department of Health Technology, Technical University of Denmark, Kongens Lyngby, Denmark; Danish Research Centre for Magnetic Resonance, Department of Radiology and Nuclear Medicine, Copenhagen University Hospital Amager and Hvidovre, Hvidovre, Denmark; Department of Neurology, Universitätsmedizin Greifswald, Greifswald, Germany; Berlin Institute of Health (BIH), 10187 Berlin, Germany; Charité – Universitätsmedizin Berlin, Humboldt-Universität zu Berlin, Berlin Institute of Health, Institute of Biometry and Clinical Epidemiology, 10117 Berlin, Germany; Neurocomputation and Neuroimaging Unit, Freie Universität Berlin, Berlin, Germany; Berlin School of Mind and Brain, Humboldt-Universität zu Berlin, Berlin, Germany; Department of Psychology, University of Greifswald, Greifswald, Germany; Research Group Cognition and Plasticity, Max Planck Institute for Human Cognitive and Brain Sciences, Leipzig, Germany; Wilhelm Wundt Institute for Psychology, Leipzig University, Leipzig, Germany; Faculty of Psychology, Chair of Lifespan Developmental Neuroscience, TU Dresden, Dresden, Germany; Centre for Tactile Internet With Human-in-the-Loop, TU Dresden, Dresden, Germany; Leibniz Research Center for Working Environment and Human Factors, Dortmund, Germany; Bielefeld University, University Hospital OWL, Protestant Hospital of Bethel Foundation, University Clinic of Psychiatry and Psychotherapy, Bielefeld, Germany; German Center for Mental Health (DZPG), Bochum, Germany; Department of Neurology, University Hospital Essen, University of Duisburg-Essen, Essen, Germany; Center for Translational Neuro- & Behavioral Sciences (C-TNBS), University Duisburg Essen, Essen, Germany; German Centre for Neurodegenerative Diseases (DZNE) Standort Greifswald, Greifswald, Germany

**Keywords:** Non-invasive brain stimulation, dose-response, structural imaging, simulation of electric fields, computational modelling, cognitive enhancement

## Abstract

Non-invasive brain stimulation is an established tool for modulating neural activity that holds promise for advancing both cognitive neuroscience and clinical interventions. Focal transcranial direct current stimulation (tDCS), using center-surround electrode montages, enables more region-specific targeting. Although computational models can simulate individual electric fields, no existing approach enables the prospective individualization of electrode placement while standardizing the dose across targeted brain regions. In the current preparatory methodological study, we present a modeling-based framework that harmonizes the electric field strength between different target regions at the group level, but preserves inter-individual variability. This enables systematic examination of dose-response relationships and their regional differences. Positioning of the center-surround electrode montages is individualized to ensure focusing of the electric field on the target regions. We started by defining brain targets for eight cognitive and motor functions using MRI data from 43 participants. Using field simulations, we then estimated a group-average field strength in the target regions that had led to behavioral and physiological effects in prior tDCS studies (resulting in 0.2 V/m). The radii of the center-surround montages were optimized for each target region to achieve the intended field strength while maximizing focality. Validation in an independent sample (n=53) confirmed that the intended target field strength is achieved on average for new participants. The described computational tools are made available as open-source software, allowing other researchers to apply our individualization framework with parameters (target regions and target field strengths) tailored to their specific research questions; and are currently being implemented in a multi-center study involving approximately 1,000 datasets.

## 1. INTRODUCTION

Non-invasive brain stimulation techniques such as transcranial direct current stimulation (tDCS) modulate neural excitability and plasticity in target brain regions associated with specific cognitive, motor, or perceptual processes (Polanía et al., 2018). Pronounced heterogeneity in tDCS outcomes highlights the urgent need to better understand and subsequently address the factors that contribute to this variability. While methodological differences across studies account for some of the observed inconsistencies, individual differences in head and brain anatomy are fundamental as they determine the amount of current reaching the brain (Polanía et al., 2018; Thielscher et al., 2011). Here, computational modeling approaches allow individualized simulations of electric fields induced by tDCS (e.g., Datta et al., 2009; Thielscher et al., 2015). Electric fields in task-relevant areas in the human brain may be closely tied to the induced behavioral and neural modulation, therefore suggesting a cortical dose-response relationship. While there have been numerous studies comparing different electrode montages, current strengths, and current densities under the electrodes as proxies for field strength in the past (Ehrhardt et al., 2021; Ghasemian-Shirvan et al., 2022; Sehm et al., 2013; Van Hoornweder et al., 2025; Zhao et al., 2024), only a few previous studies have investigated potential associations between electric field strengths induced at the individual level and the corresponding modulation of functional brain connectivity and behavioral performance (Antonenko et al., 2017; Indahlastari et al., 2021; Jamil et al., 2020; Kim et al., 2014). For example, higher electric fields resulted in more pronounced enhancement of verbal working memory (Albizu et al., 2020; Kim et al., 2014) or functional connectivity changes in the stimulated brain network (Antonenko et al., 2019; Indahlastari et al., 2021). Moreover, cortical electric field strength correlated with neurochemical and functional connectivity changes in the sensorimotor network (Antonenko et al., 2019; Nandi et al., 2022). A recent study concluded that regional anatomical features (such as electrode-to-cortex distance) together with electric field strength predict local neurophysiological tDCS effects on motor evoked potentials and cerebral blood flow (Mosayebi-Samani et al., 2021). Such evidence stems from small exploratory studies and only specific brain networks have been investigated. Further, there are also studies which have not observed clear associations of simulated electric fields and interventional outcomes (Liu et al., 2022; Mezger et al., 2021; Muffel et al., 2019; Nandi et al., 2022). Thus, to date, the relationship between cortical electric field dose of tDCS and the behavioral and neurophysiological stimulation outcomes as well as the factors influencing this relationship (such as age or underlying brain microstructure), are not well understood. In addition, it is unclear how much these relationships differ across brain regions and functional domains.

To address these open questions, we aimed to develop a dosing and targeting approach for focal tDCS using center-surround montages to allow for more region-specific targeting of brain regions compared to conventional tDCS montages (Alam et al., 2016; Kuo et al., 2013; Niemann, Shahbabaie, et al., 2024). Compared to conventional electrode montages, focal tDCS montages may be better suited to study dose-response relationships for cortical areas, as they minimize the unintended co-stimulation of areas surrounding the target sites. Our method is thus specifically designed for center-surround configurations, as their better spatial specificity of stimulation facilitates the interpretation of dose-response relationships in functional magnetic resonance imaging (fMRI) studies. Specifically, it becomes more straightforward to distinguish local effects (within the targeted region) from remote network effects.

We opted for an approach that (1) harmonizes the average electric field dose across different target regions, (2) allows prospectively usages, i.e., to plan electrode configurations before focal tDCS is applied in a participant, and (3) maintains interindividual dose variability in each target to enable systematic tests of dose-response relationships. First, to define the physiologically relevant target electric field, we employed electric field simulations in a preparatory computational modeling study: As there is currently no general consensus regarding the minimal current intensity threshold required for physiological modulation and no knowledge about the potential optimization criteria for effective delivery (Fertonani & Miniussi, 2017; Lee et al., 2021), we decided to define the target electric field magnitude as the averaged field strength across individuals and domains, induced by previous electrode montages for tDCS that have reported beneficial behavioral effects, using individualized field simulations in Sample 1 (n=43 healthy participants, age range from 20-47 years). The strategy is generic and can be applied beyond this multi-center study.

We then systematically varied the center-surround radii of the simulated focal tDCS montages to determine project-specific radii that on the group-level resulted in a similar electric field strength in the target regions (Dmochowski et al., 2011; Radecke et al., 2020; Saturnino et al., 2020; Saturnino, Siebner, et al., 2019). To ensure region-specific spatial targeting, the center electrodes were thereby individually placed at skin positions with minimal distance to the centers of the target regions (minimizing “unwanted” co-stimulation, thus robustly constraining the current on the intended target area only). Simulations in a second, independent cohort (Sample 2: n=53 healthy participants, age range from 19-42 years) were used to confirm that the chosen region-specific radii achieved similar group-level field strengths in the target regions also for new participant groups. Additionally, a control experiment showed that the placement strategy for the center electrode position based on Euclidean distances performed well. It compared favorably when to a more principled positioning approach that accounts for tissue conductivities beneath the anode to ensure optimal electrical connection of the anode to the ROI.

In the current methodological paper, we delineate the process of determining the target regions and the target field strength using previous evidence and actual simulations of electric fields across two study samples, followed by the development of an algorithm for automatic individualized placement of the electrode positions of the focal tDCS montages (see **Figure 1** for a visual illustration of the study design).

**Figure 1.**
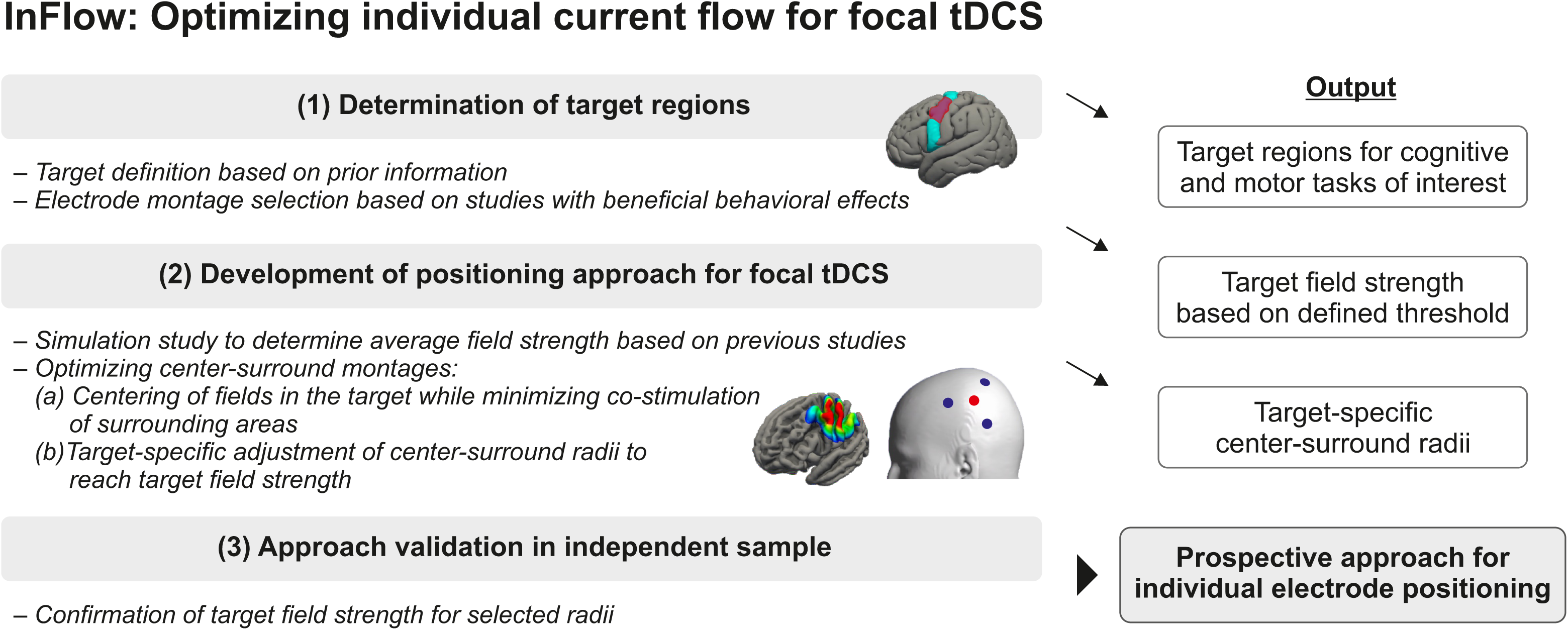
Illustration of the study design. First, we determined target regions based on prior information about location and spatial extent of target regions mediating the respective cognitive or motor function. This information was derived from previous fMRI experiments using similar tasks as those to be implemented in the multi-center study. We then selected studies using electrode montages for tDCS that have reported beneficial behavioral effects in similar functions. This can be done for any target regions of interest, depending on the cognitive or motor domain or the task under study. Second, to define the physiologically relevant target electric field, we employed electric field simulations in a preparatory computational modeling study in Sample 1 (n=43, 20-47 years). These analyses resulted in an average behaviorally relevant electric field strength of 0.2 V/m for the electrode montages employed in the selected studies. This results in a target field strength which is specific to our multicenter study and can be defined based on other decisions or approaches. Third, switching to focal center-surround tDCS montages, we systematically varied the center-surround radii in further simulations to determine target-specific radii that resulted in the desired electric field strength of 0.2 V/m in the target region while minimizing co-stimulation of surrounding areas. The electric field intensity in the brain region under the center electrode generally increases with the radius, as this reduces the amount of current shunted through the skin. Please note that our aim was to select project-specific (rather than personalized) radii so that the desired electric field strength of 0.2 V/m was reached on average across the group. Fourth, simulations in an independent cohort (Sample 2: n=53, 19-42 years) were used to confirm the selected region-specific radii. This prospective approach for individual electrode positioning can be implemented for personalization based on individual MR scans, or for designing electrode montages that produce comparable field strength across different brain regions.

Our approach is available as open-source code (https://github.com/simnibs/memoslap_utils). It can be implemented for individual electrode positioning for personalization based on individual MR scans, or for designing electrode montages that produce comparable field strength across different brain regions. With the input of T1- and T2-images for each individual and the target region, the algorithm generates a single report file containing coordinates for a focal electrode montage for use in neuronavigation systems.

## 2. METHODS

### 2.1. Participants

Structural brain images of 43 healthy adult participants (23 females, age range: 20-47 years, mean/SD education: 16.6/2.1, all right-handed) were acquired (Sample 1) to test our approach. To ensure reproducibility of estimated field magnitudes and focalities, we added a re-test dataset of 53 participants (26 females, age range: 19-42 years; Sample 2, mean/SD education: 15.8/2.5, all right-handed). MR images of Sample 1 participants were acquired in the context of a study examining electrophysiological memory correlates and modulation. MR images of Sample 2 participants were acquired in the context of the multi-center project for which we aimed to develop the method. Exclusion criteria for participation in both studies were a history of neurological or psychiatric diseases, metal or electronic implants (e.g., pacemakers, surgical devices), and pregnancy. The studies were approved by the ethics committee of the Greifswald University Medicine and conducted in accordance with the Helsinki Declaration (Sample 1: BB 021/19; Sample 2: BB 015/22, 016/22, 017/22). Written informed consent was obtained from all participants prior to participation.

### 2.2. Magnetic resonance imaging

Sample 1 data were acquired on a 3T Siemens Verio equipped with a 32-channel head coil (Institute of Radiology, University Medicine Greifswald, Germany). High-resolution T1-weighted (1x1 x1 mm³, TR = 2300 ms, TE = 2.96 ms, TI = 900 ms, flip angle = 9°; using selective water excitation for fat suppression) and T2-weighted images (1x1x1 mm^3^, TR = 12770 ms, TE = 86 ms, flip angle = 111°) were recorded. Sample 2 data were acquired on a 3T Siemens Vida using a 64-channel head coil, high-resolution T1- (0.9x0.9x0.9 mm³, TR = 2700 ms, TE = 3.7 ms, TI = 1090 ms, flip angle = 9°; using selective water excitation for fat suppression) and T2-weighted images (0.9x0.9x0.9 mm³, TR = 2500 ms, TE = 349 ms) were recorded.

### 2.3. Determination of target regions

To identify target regions, we employed an iterative approach that considered both anatomical landmarks, previous task-related fMRI and transcranial direct current stimulation (tDCS) studies that have investigated the respective cognitive or motor function targeted in the eight projects (P1-8) of the multi-center study: P1, object-location learning; P2, tacto-spatial working memory; P3, novel-word learning; P4, verbal working memory; P5, motor sequence learning; P6, eyeblink conditioning; P7, learning-based control; P8, value-based learning. Atlas-based brain areas (Desikan et al., 2006) showing the highest overlap with the regions targeted in the empirical projects were determined as regions-of-interest (ROIs): P1, right occipitotemporal cortex (Gillis et al., 2016), P2, left superior parietal lobe (Schmidt et al., 2021; Wang et al., 2019), P3, left inferior frontal gyrus/pars opercularis (Filippova et al., 2022; Perceval et al., 2020; Perikova et al., 2022), P4, left inferior frontal gyrus/pars opercularis (Emch et al., 2019; Niu et al., 2019), P5, left primary motor/precentral cortex (Nitsche et al., 2003; Pimentel et al., 2013), P6, right cerebellar lobe (corresponding to Crus I and Lobule VI from the SUIT atlas (Diedrichsen & Zotow, 2015)) (Zuchowski et al., 2014), P7, right dorsolateral prefrontal/rostral middle frontal cortex (Gbadeyan et al., 2016; Kerns et al., 2004), P8, left dorsolateral prefrontal/rostral middle frontal cortex (Eppinger et al., 2015; Nikolin et al., 2018). All ROIs except the right cerebellar lobe (P6) were defined on fsaverage space, to leverage the higher accuracy of surface-based (using CAT12 (Gaser et al., 2024) functionality embedded in SimNIBS charm, see below) compared to volume-based registrations when transforming ROIs to the individual brain (Duecker et al., 2014). The ROI of P6 was determined in MNI space. The resultant group-level ROIs were created by overlapping the atlas regions with spheres of 20-mm radius around the center of gravities of fMRI activity (except for P6 where the atlas-based ROI was used given its specific parcellation) to achieve more regional precision and corresponded to parts of the gyrus relevant for the respective cognitive/motor function. Group-level ROIs were then mapped on each individual brain to obtain individual regions. For P1-P5, P7, P8, the group-level ROIs were mapped to cortical middle gray matter sheet in individual space using the surface-based transformation functionality in SimNIBS. For P6, non-linear volume-based transformation from SimNIBS was used to transform the ROI from MNI into individual space, where it was then intersected with the cerebellar middle gray matter surface (estimated as described below).

### 2.4. Calculation of ROI sizes

For calculating individual ROI sizes, the group-level ROIs of the left and right hemispheres of the cerebrum were mapped to the individual middle gray matter surfaces, applying the surface-based registration of CAT12 embedded in SimNIBS. The group-level ROI of P6 (in the right cerebellar lobe) was non-linearly transformed from MNI to individual space using SimNIBS functionality and intersected with the cerebellar middle gray matter surface. The ROI size (in [mm²]) was then determined using SimNIBS functionality.

### 2.5. Head segmentation

Tissue segmentation of the head and brain from T1- and T2-weighted images was performed using charm from SimNIBS version 4.0 (simnibs.org) (Puonti et al., 2020; Thielscher et al., 2011). The generated individual tetrahedral head meshes including representations of the scalp, skull, spongy bone, compact bone, cerebrospinal fluid (CSF), gray matter, white matter, large blood vessels and eyes.

To enable a consistent surface-based analysis approach for all projects, a surface that approximates the middle of the cerebellar gray matter was reconstructed from the volume segmentation results obtained by charm. This was done by assigning different isopotential values (0.0 and 1.0) to the boundaries between the segmented cerebellar gray matter and CSF, as well as between cerebellar gray and white matter. The Laplace equation was then solved to determine the isopotential surface for 0.5, indicating the middle positions between the two boundaries. Owing the limited image resolution of clinical structural MR images, the fine cortical folding structure of the cerebellum is not resolved in the volume segmentation and also not present in the reconstructed middle surfaces. The electric field estimates in the cerebellum thus only capture the impact of the gross cerebellar anatomy on the induced field. Individual datasets were visually inspected in order to assure accurate head reconstructions (Nielsen et al., 2018). All data were deemed appropriate.

### 2.6. Calculation of tissue volumes

The volumes of CSF, skull (compact and spongy bone combined) and scalp were extracted in cylindrical volumes of interest underneath the center electrodes of the focal tDCS montages for cortical targets. The cylinders were oriented orthogonally to the local skin surface extracted from the tetrahedral head meshes, had radii of 20 mm and reached 40 mm into the head mesh. The tissue volumes in mm³ were calculated by summing the volumes of the tissue-specific tetrahedra of the head mesh inside the cylinders.

### 2.7. Simulation of electric fields for “empirical” electrode montages to determine the target field strength (Sample 1)

Stimulation parameters were defined in SimNIBS corresponding to the actual setups in previous studies targeting the respective functional domains (**Supplementary Figure 1** for illustration of electrode positions and **Supplementary Table 1** for project and task details and **Supplementary Table 2** for stimulation parameters). Median electric field strength (in V/m), averaged within the target ROI and field focality (quantified as area of field values above the median) were extracted from individual grey matter central surface outputs. Field strength (i.e., the vector norm of the electric field in V/m) within the ROIs was defined as target variable due to its independence from direction of the field and based on previous evidence for associations with anatomical features and empirical effects (Saturnino, Puonti, et al., 2019). The median field strength across all ROIs was used as target field strength for the subsequent optimization of the focal montages. Stimulation of electric fields for focal montages were performed using four round electrode (of 2 cm diameter) and a current intensity of 2 mA.

### 2.8. Position approach for focal tDCS

Montage individualization aims to ensure that the fields induced by the focal montages are robustly centered on the individual ROIs. Our approach consists of two parts: First, placing the anode above the ROI center to optimize the focality-intensity trade-off of the induced electric fields. Second, reducing the radius of the montage as much as possible to minimize the co-stimulation of other areas while ensuring the desired field strength in the target. Of note, optimization of the radii in the second step was done on the group level, with the aim to achieve the same field strength in the different target regions on average across participants while preserving inter-individual variability. Details for these two steps are given below. See also **Supplementary Material B** for further analyses to evaluate the performance of our approach.

We developed and compared two algorithms that systematically adjust the position of the anode. Algorithm 1 (termed “Euclidean optimization”) minimizes the Euclidean distance of the anode position on the scalp surface to the ROI centers. However, considering the complex head anatomy, the smallest Euclidean distance might not correspond to the scalp position that is electrically “best” connected to the ROI. Thus, algorithm 2 (“resistivity optimization”) instead solves a Laplace equation using the finite element method (FEM) of SimNIBS, where the cortical ROI and the remaining cortex surface are set to different isopotential values (1.0 and 0.0). This choice corresponds to weighting all parts of the ROI equally as target and treating the rest of the brain surface homogeneously as avoidance region, which is the standard choice in “classical” optimization algorithms for multi-channel stimulation for optimizing focality (Dmochowski et al., 2017; Saturnino, Siebner, et al., 2019). The scalp position with the highest potential value is then chosen as anode position. This ensures that the electric resistance between the anode position and the ROI is chosen as small as possible relative to the resistance between the anode position and the rest of the cortex. Both algorithms performed very similarly (see Results), confirming the suitability of algorithm 1 that was finally chosen as it was computationally more efficient and faster in practice.

The three surround electrodes were distributed at equal angles around the center. We chose three instead of four cathodes due to practical feasibility. Note, however, that using more than three cathodes does not markedly change the field (Videira et al., 2022). We replicated this previous result in our analyses, showing nearly identical field magnitudes and focality in 4x1 and 3x1 montages (see **Supplementary Figure 2** for an illustration of magnitude comparability). Orientation was varied for different angles (between 0 and 105° in 15°-steps). As the orientation of the cathodes did not systematically impact electric field magnitudes on the group level (**Supplementary Figure 3**), we selected a practically feasible phi-offset for each project (for instance, not covering the ears or eyes). Center-surround radii between the anode and the three surrounding cathodes were varied from 40 to 75 mm and simulated for each subject in 5 mm steps, employing a fixed current intensity of 2 mA. For each ROI, the minimal center-surround radius was selected which ensured that the desired target field strength was on average reached while keeping the field as focal as possible.

### 2.9. Statistical analyses

R was used for statistical analyses (R Core Team, 2020). Center-surround radii were determined on Sample 1 data while all further statistical comparisons were calculated on Sample 2 data. For each dependent variable, we conducted Cohen’s d to quantify the effect size of pairwise comparisons between projects, based on a linear mixed model approach. Cohen’s d is a standardized metric that expresses the magnitude of the difference between two groups, accounting for the variability within each group. To calculate Cohen’s d for each pairwise project comparison, we first estimated the model based marginal mean difference between the groups and then divided it by the pooled standard deviation of the measurements. This method allowed us to assess the observed differences between project performances, considering both the effect size and the variability inherent in the data in a standardized way. Linear mixed model analyses (random intercept models) were calculated for each dependent variable (magnitude or focality). Independent variables were the project (project 1 to 8), algorithm (1 or 2), age, and sex, with an interaction term between project and algorithm included to assess potential variations in the effect of algorithm across different projects. A random intercept term was included to account for the repeated measurements or nested structure of the data, where a subject represents the individual units of observation. To further investigate pairwise comparisons of project variances, we calculated the log-transformed coefficient of variation ratio (CVR) for each pair of projects. CVR was computed as the ratio of the coefficient of variation (CV) of the magnitude and focality between two projects.

## 3. RESULTS

### 3.1. Target regions

Target regions delineated based on anatomical information as well as functional magnetic resonance imaging (fMRI) and tDCS studies are illustrated in **Figure 2A**. The eight projects (P1-8) of the multi-center study were: P1, object-location learning; P2, tacto-spatial working memory; P3, novel-word learning; P4, verbal working memory; P5, motor sequence learning; P6, eyeblink conditioning; P7, learning-based control; P8, value-based learning. We observed interindividual variability as well as differences between projects in ROI sizes with low values in project 1 (Cohen’s d between 0.71 and 2.36; compared with projects 2, 3, 4, 5, 7, 8) and project 6 (Cohen’s d between 1.01 and 2.6; compared with projects 2, 3, 4, 5, 7, 8) and high values for project 2 (Cohen’s d between 0.86 and 2.6; compared with projects 1, 4, 5, 6, 7, 8) and high values in project 3 (Cohen’s d between 0.72 and 1.58; compared with projects 1, 2, 5, 6; **Supplementary Figure 4**). Although ROI sizes were not associated with electric field magnitudes (overall r = 0.25, p = 0.797) or focality from the focal tDCS simulation (overall r = 0.172, all p = 0.862), subsequent group-level analyses were adjusted for ROI sized to account for differences. To further explore how potential head and brain anatomical differences between the target regions can affect electric fields, we compared different tissue volumes (i.e., skin, skull, CSF) below the electrodes. Volumes differed between cortical targets for skin (Cohen’s d’s between 0.01 and 2.31) and skull (Cohen’s d between 0.01 and 2.20). These analyses revealed that mainly CSF volumes were lower for the occipitotemporal cortex (target in P1) compared to the other target regions (Cohen’s d: 0.91-3.13), confirming that anatomical factors that differ systematically may indeed affect the comparability between regions (**Supplementary Figure 5**). Larger CSF volume was associated with lower electric field magnitudes at target areas (r_rm_ = -0.72, p < 0.001; **Supplementary Figure 6**) and explained 70 % of their variance (adjusted for skin and skull volume).

**Figure 2.**
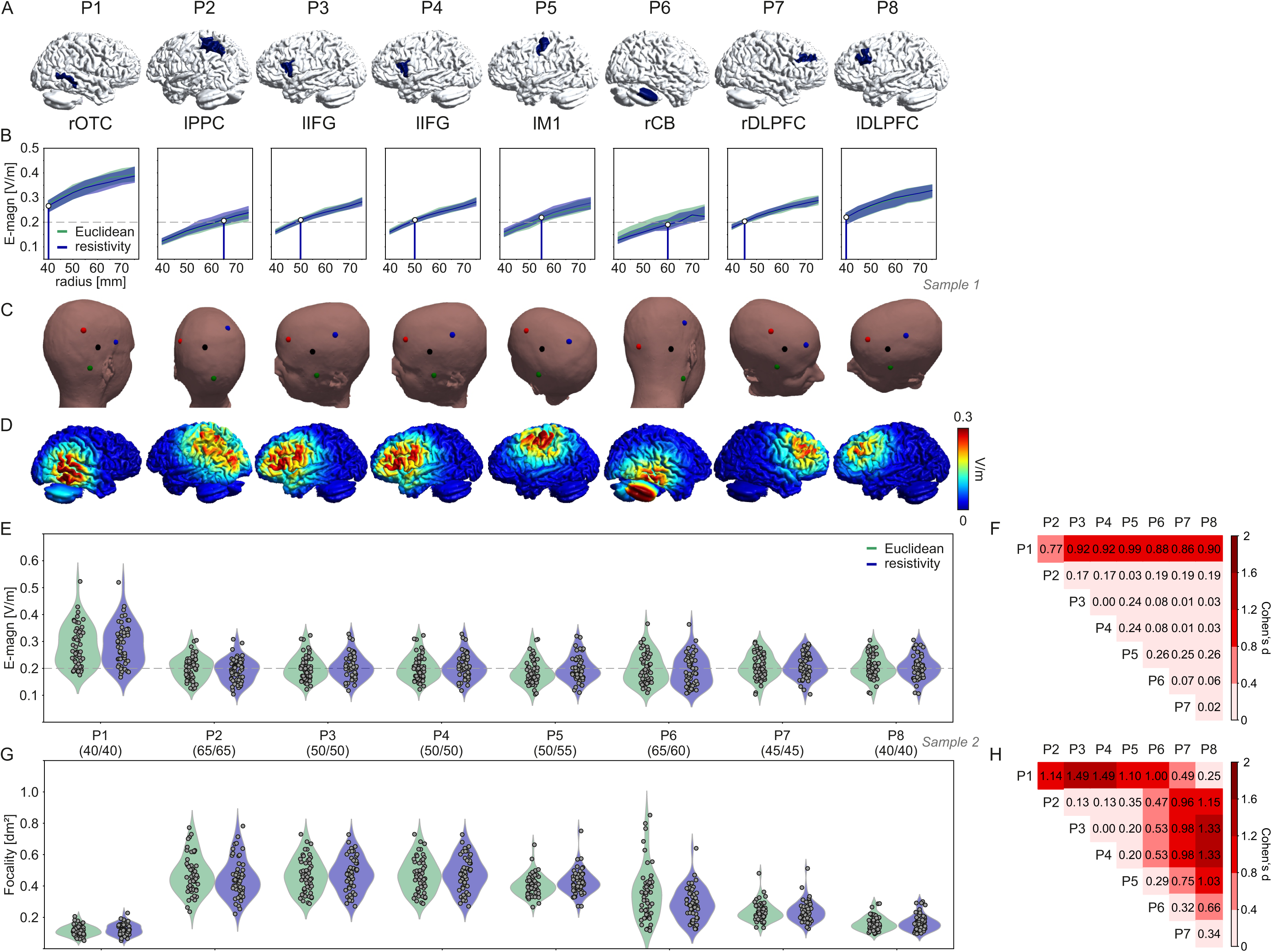
(A) Target regions for individual projects of the research unit “MeMoSLAP” (eight projects P1-8: P1, object-location learning; P2, tacto-spatial working memory; P3, novel-word learning; P4, verbal working memory; P5, motor sequence learning; P6, eyeblink conditioning; P7, learning-based control; P8, value-based learning) illustrated for one subject from the sample. (B) Variation of distance between center anode and the three surround cathodes (Sample 1, n=43). The dashed line represents the average electric field strength (0.2 V/m) that resulted from preparatory computational modeling analyses of “empirical” montages yielding beneficial tDCS effects. (C) Resultant electrode positions of the 3x1 focal tDCS set-ups for focal tDCS in P1-8 with matched average electric field magnitudes in target regions for the “Euclidean” algorithm. Note: while positions for the anodes were optimized for individual subjects, residual variability within studies will allow for testing cortical dose-relationships. (D) Sample e-field distributions for the given 3x1 set-ups. (E) Average electric field magnitude induced in target ROIs in 53 participants (Sample 2) with the resultant set-up for P1-8 (radii of Euclidean/resistivity algorithms in brackets). (F) Pairwise comparison of electric field magnitude between projects (Cohen’s d; n=53 subjects, number of obs: 848). (G) Focality of tDCS induced in target ROIs in 53 participants with the resultant set-up for P1-8. (H) Pairwise comparison of electric field focality between projects (Cohen’s d; n=53 subjects, number of obs: 848). rOTC, right occipitotemporal cortex. lPPC, left posterior parietal cortex. lTP, left temporoparietal. lIFG, left inferior frontal gyrus. lM1, left motor cortex. rCB, right cerebellum. rDLPFC/lDLPFC, right/left dorsolateral prefrontal cortex.

### 3.2. Target field strength

Using data of Sample 1, we simulated the field distribution for eight “empirical montages”. We observed high variability of the average median electric field strength in the target ROIs across participants, with a range from 0.06 to 0.39 V/m across electrode montages (median (IQR) P1: 0.14 (0.03), P2: 0.28 (0.03), P3: 0.13 (0.02), P4: 0.25 (0.04), P5: 0.31 (0.05), P6: 0.21 (0.04), P7: 0.11 (0.03), P8: 0.14 (0.05); **Supplementary Figure 7A**) with large differences between projects (**Supplementary Figure 7B**). The overall average across projects was 0.20 V/m.

Focality values ranged between 0.10 to 9.96 dm² (median (IQR) P1: 2.29 (1.99), P2: 3.31 (1.19), P3: 0.65 (0.13), P4: 2.80 (0.70), P5: 1.91 (0.77), P6: 0.48 (0.41), P7: 0.27 (0.10), P8: 0.19 (0.09); **Supplementary Figure 7C**) with likewise large differences between projects (**Supplementary Figure 7D**).

Based on these simulations, we defined a field strength of 0.2 V/m as the “target field strength” to be reached over the study sample on average for each of the projects. Note, that this threshold resulted from averaging field magnitudes in all positions within the target region, which is lower than the peak dose. As substantial parts within the target areas exceed the threshold of 0.2 V/m, we additionally computed the 95th percentile in the target regions (Mirjalili et al., 2025). The results (**Supplementary Figure 8**) are in line with previously reported “effective doses” of 0.3 V/m (SD 0.11 V/m) for tACS (Alekseichuk et al., 2022) and for distinguishing responders from non-responders to tDCS (Mirjalili et al., 2025).

### 3.3. Group-level optimization of center-surround radii for focal brain stimulation

Electric field magnitudes of Sample 2, averaged within the target ROIs, varied in dependence of the radius between the center anode and surrounding cathodes (**Fig. 2B**) with higher magnitudes for larger distances. The two algorithms resulted in almost identical field magnitude estimations, warranting the “Euclidean optimization” as valid. The center-surround distances (optimized for each ROI) required for a median field strength of 0.20 V/m varied between 40 and 65 mm across the eight target ROIs (**Fig. 2C** for electrode placements on a sample head, **Fig. 2D** for sample field distributions). Please note that the median field strength of P1 stayed higher than the desired target field strength even for the smallest radius of 40 mm. We did not reduce the radius further in order to maintain practical feasibility and prevent unintended shunting by electrode gel bridges. Variation of the phi offset did not affect electric field magnitudes (**Supplementary Fig. 3**). Therefore, we selected a practically feasible phi-offset for each project (for instance, not covering the ears or eyes).

#### 3.3.1. Magnitude

Individual electric field magnitudes within the target regions were all distributed around the target field strength of 0.2 V/m, except the occipitotemporal cortex (P1) where the group-based average lay above this threshold. Accordingly, a statistical comparison of electric field magnitudes revealed differences between projects with higher values for P1 compared to the others (all Cohen’s d ≥ 0.77, for all other pairwise comparisons Cohen’s d ≤ 0.24; **Fig. 2E**, **Fig. 2F**).

Coefficient of variation (CV) of electric field magnitudes, which quantifies their spread relative to their mean and thus represents inter-individual variability, showed higher variability in P6 and lower variability in P1 compared to the other projects (**Supplementary Figure 9A**).

#### 3.3.2. Focality

Focality of the induced electric fields differed between projects with substantially higher focality (corresponding to lower values) in P1 (range of Cohen’s d: 1.00-1.49; compared with P2, P3, P4, P5, and P6), P7 (range of Cohen’s d: 0.75-0.98; compared with P2, P3, P4, and P5) and P8 (range of Cohen’s d: 0.66-1.33; compared with P2, P3, P4, P5, P6; **Fig. 2G**, **Fig. 2H**). Coefficient of variation of electric field focality showed some differences between projects (**Supplementary Figure 9B**).

## 4. DISCUSSION

The current study presents the methodological development and validation of a positioning approach for focal tDCS that harmonizes the stimulation dose across target sites using electric field simulations. This includes the estimation of an “effective” dose from previous studies, algorithm development for electrode positioning, determination of optimal region-specific center-surround radii of the focal montages, and validation in an independent sample. Our work presents a preparatory study to allow prospective region-specific positioning of electrodes in studies investigating the relationship between electric field dose and the resulting activity and performance modulation. Our open-source tools (https://github.com/simnibs/memoslap_utils) allow researchers to easily tailor montages to their intended target regions and target field strength.

### 4.1. Target field strength

Currently, there are no established, evidence-based, region-specific electric field thresholds for different brain areas. No general consensus exists regarding the minimal current intensity threshold required for physiological modulation which may even differ between individuals (Fertonani & Miniussi, 2017; Van Hoornweder et al., 2025). Studies have shown that higher electric fields (e.g., 0.25 V/m in left DLPFC) can produce stronger working memory improvements for tDCS (Caulfield et al., 2022). Some studies have reported thresholds around 0.2 to 0.3 V/m for distinguishing tDCS responders from non-responders (Mirjalili et al., 2025). Similar field strength thresholds have been reported by Alekseichuk et al. (2022) for inducing effects in awake/behaving mammals by tACS (mean ± SD across five studies: 0.23 ± 0.10 V/m). In contrast, the required field strengths in brain slices and anesthetized rodents seem to be generally higher (Alekseichuk et al., 2022; Fritsch et al., 2010). Together, previous evidence supports that our target of 0.2 V/m is within a physiologically relevant range (Mirjalili et al., 2025; Van Hoornweder et al., 2025). However, optimal thresholds for specific brain regions remain undetermined.

Our approach used a uniform average “effective” target field strength across all target regions. We followed the rationale that real tDCS applied in previous studies, yielding beneficial effects for 1-2 mA, would inform the selection of an average electric field magnitude needed to be achieved. This is a pragmatic, evidence-based starting point that accounts for interregional differences in head and brain anatomy that systematically affect the induced field. However, several local factors including cortical network organization and neuron properties including cytoarchitecture are likely to affect the physiological response thresholds. Thus, by harmonizing the group-average field strength across target regions while keeping the interindividual variability of the field strength, our goal here was to provide the methodological basis for systematic comparisons of dose-response relationships across brain areas and related cognitive and motor domains. This systematic evaluation and comparison of minimal effective doses across regions is possible as our approach results in different electric field strengths in each participant depending on head and brain anatomy.

Importantly, the provided open-source code allows adaptation of the “desired” region-specific field strength. Our dedicated aim was to ensure that the montages induce similar, behaviorally effective electric field ranges in different target regions while being as focal as possible.

### 4.2. Precision stimulation

A reliable method for prospective planning of focal tDCS application considering optimized individual spatial targeting is currently not available. Electrode placements in focal tDCS most often rely on the 10-20 electroencephalographic system. Due to interindividual anatomical differences, this approach may result in inaccurate electrode positioning in individual participants, thereby introducing variable electric fields in the target region (Huang et al., 2017; Niemann, Riemann, et al., 2024; Opitz et al., 2018; Woods et al., 2015). Our approach ensures that the focal tDCS montages are accurately centered above the target regions, with neuronavigation being used to ensure that the planned electrode positions are reliable reached in practice (Niemann, Shahbabaie, et al., 2024). In addition, our approach allows validating electrode placement based on a simple minimization of the Euclidean distance to the brain target, using a more principled approach that accounts for the impact of the different tissue conductance on the current pathways in the head.

## 0¡9¡ Limitations

For the P1 montage, the average target field strength remained above 0.2 V/m even at the lowest practically feasible center-surround radius of 40 mm. Control analyses quantifying the volumes of scalp, skull, and CSF underneath the electrodes demonstrated that local anatomical factors differed systematically between the target positions. Mainly CSF volume, which was substantially lower above the occipitotemporal cortex. These differences may limit the effectiveness of our strategy, which aims to homogenize the induced field strengths across positions by solely adjusting the center-surround radii. However, we chose not to reduce the radius below 40 mm to maintain practical feasibility and avoid shunting between electrodes. This could also impact dose-response relationships, making it more challenging to compare P1 with the other projects. While reducing the applied current intensity would have been a straightforward solution to lower the target field strength, we here opted against it to maintain a consistent current intensity across all projects, also aiming to keep the sensory side effects similar. Further, modeling of the detailed folding structure within the cerebellar ROI (P6) was lacking. The reconstructed surface is placed at half of the depth of the cerebellar gray matter in order to ensure that the average field strength in the ROI is close to the average field strength in the true, highly folded gray matter. However, as the area of the cerebellar gray matter is strongly underestimated by the surface, the reported focality values for the cerebellar ROI are too small.

As our study was part of preparatory work for a multi-center study, we selected the age range of participants based on its aims (https://osf.io/t37u2). This will allow investigating age effects on stimulation outcomes, but may limit conclusions outside this age range. Importantly, our tool can be applied for other age groups as well.

## 0¡0¡ Conclusions

We developed a strategy for estimating an effective electric field strength based on previous functional imaging and behavioral studies comparing real and sham tDCS. We used the results to inform an approach that ensures accurate spatial targeting of focal brain stimulation and enables the harmonization of the induced electric field strength across different target region. Here, we provide a methodological framework to allow prospective planning of region-specific tDCS application and enable systematic assessments of interindividual and interregional differences in tDCS dose-response profiles. Whether it proves behaviorally effective requires future studies and is currently being investigated in a multicenter study.

## Acknowledgement

This study was performed in the context of the multi-center study MeMoSLAP - “Modulation of brain networks for memory and learning by transcranial electrical brain stimulation: A systematic, lifespan approach” (Research Unit FOR 5429, funded by the German Research Foundation: Deutsche Forschungsgemeinschaft, DFG; https://osf.io/t37u2).

## Code availability

The underlying code for this study is available in GitHub and can be accessed via this link https://github.com/simnibs/memoslap_utils.

## Data availability

The datasets analysed during the current study are not publicly available due to potential identifying information that could compromise participant privacy, but are available from the corresponding author on reasonable request.

## Author contributions

AT: Conceptualization, Software, Formal analysis, Writing – original draft, Writing – review & editing, Funding acquisition. DH, UG: Formal analysis, Visualization. OP: Software, Formal analysis. FB, RF, GH, AF, SCL, MM, MN, DT: Writing – review & editing, Funding acquisition. DA: Conceptualization, Formal analysis, Visualization, Investigation, Writing – original draft, Writing – review & editing, Visualization, Funding acquisition. All authors read and approved the final manuscript.

