## Supplementary Material A for "Harmonizing the stimulation dose of focal tDCS across target sites"

### Supplementary Material A - Figures and Tables

**Supplementary Figures**

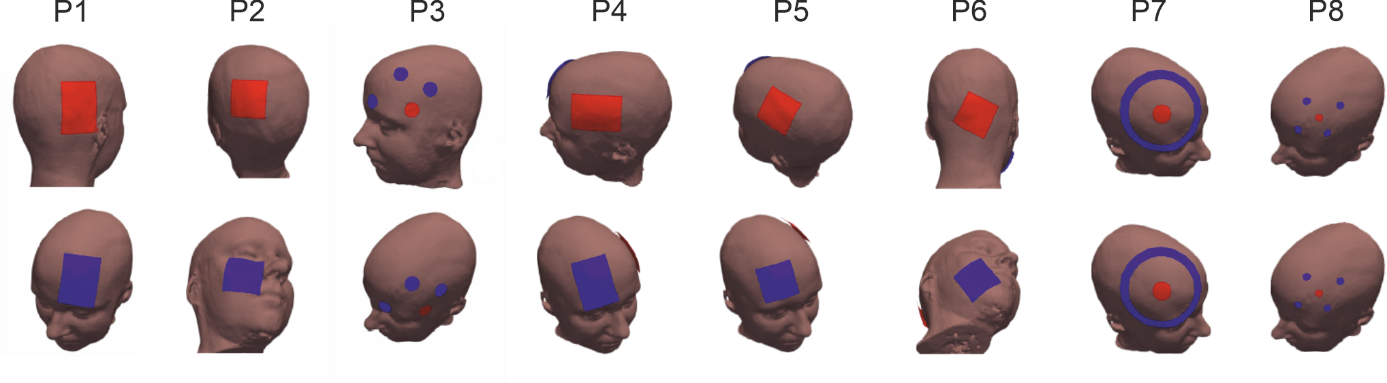

***Supplementary Figure 1.*** Electrode montages for projects 1-8 (P1-8) of the Research Unit “MeMoSLAP” yielding beneficial effects in previous studies selected from the existing evidence (termed “empirical montages”), red: anode; blue: cathode(s).

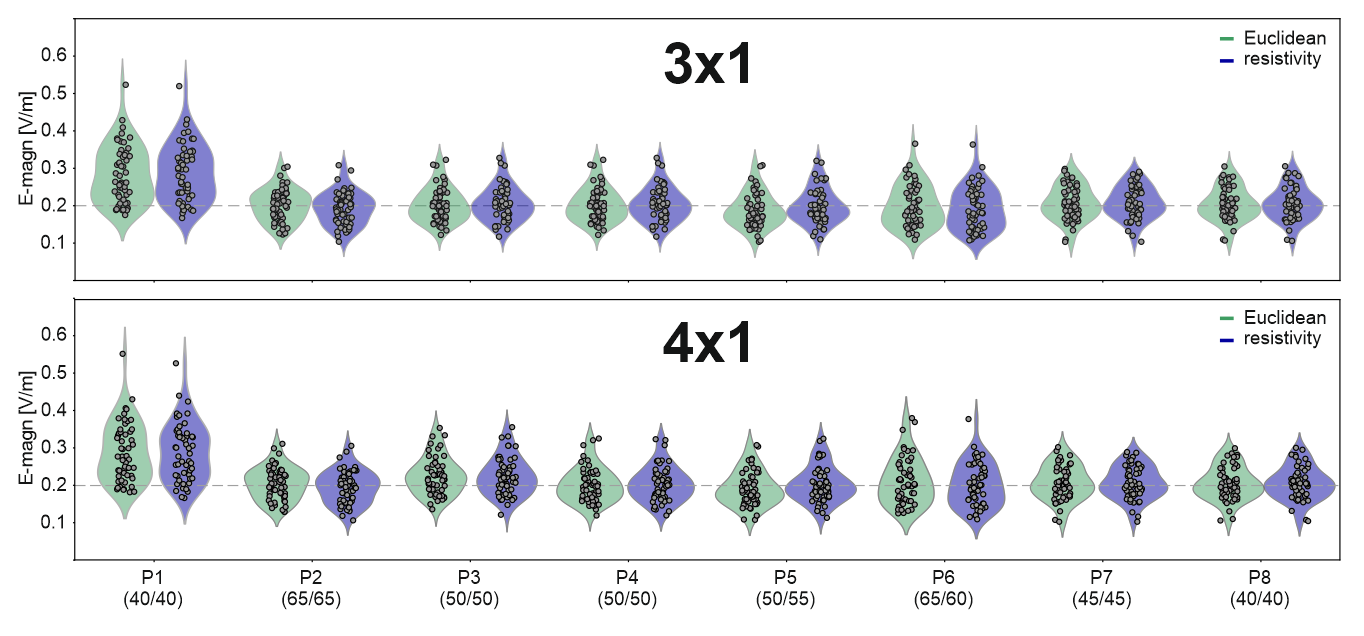

***Supplementary Figure 2*.** E-field magnitude distributions for the focal tDCS set-ups in the target ROIs of P1-8, for 3x1 (top) versus 4x1 (bottom), in 53 participants (Sample 2).

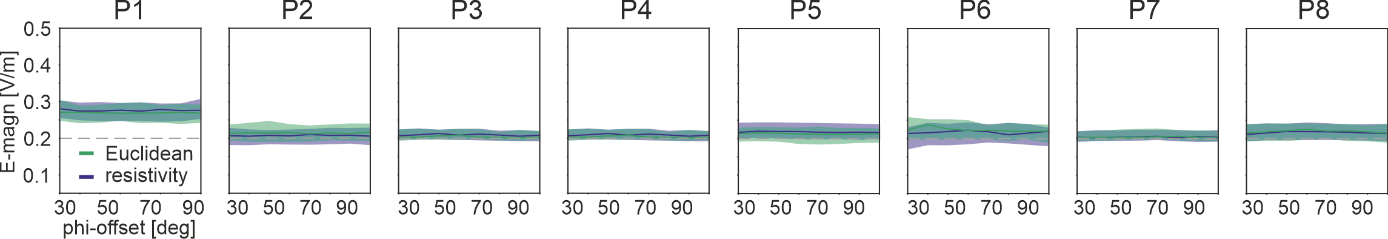

***Supplementary Figure 3.*** Variation of the orientation (i.e., phi-offset) of cathodes around the center anode (using the “best” radius determined in previous analyses).

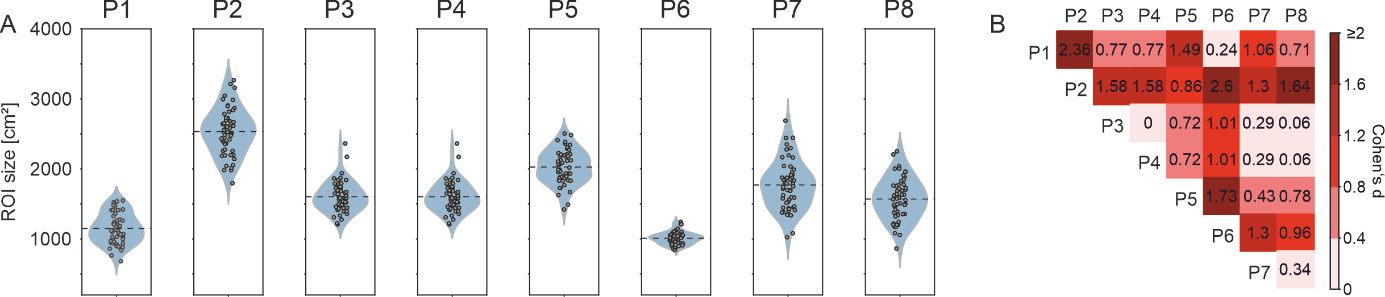

***Supplementary Figure 4.*** ROI sizes of grey matter target regions (A) and their pairwise comparison (B).

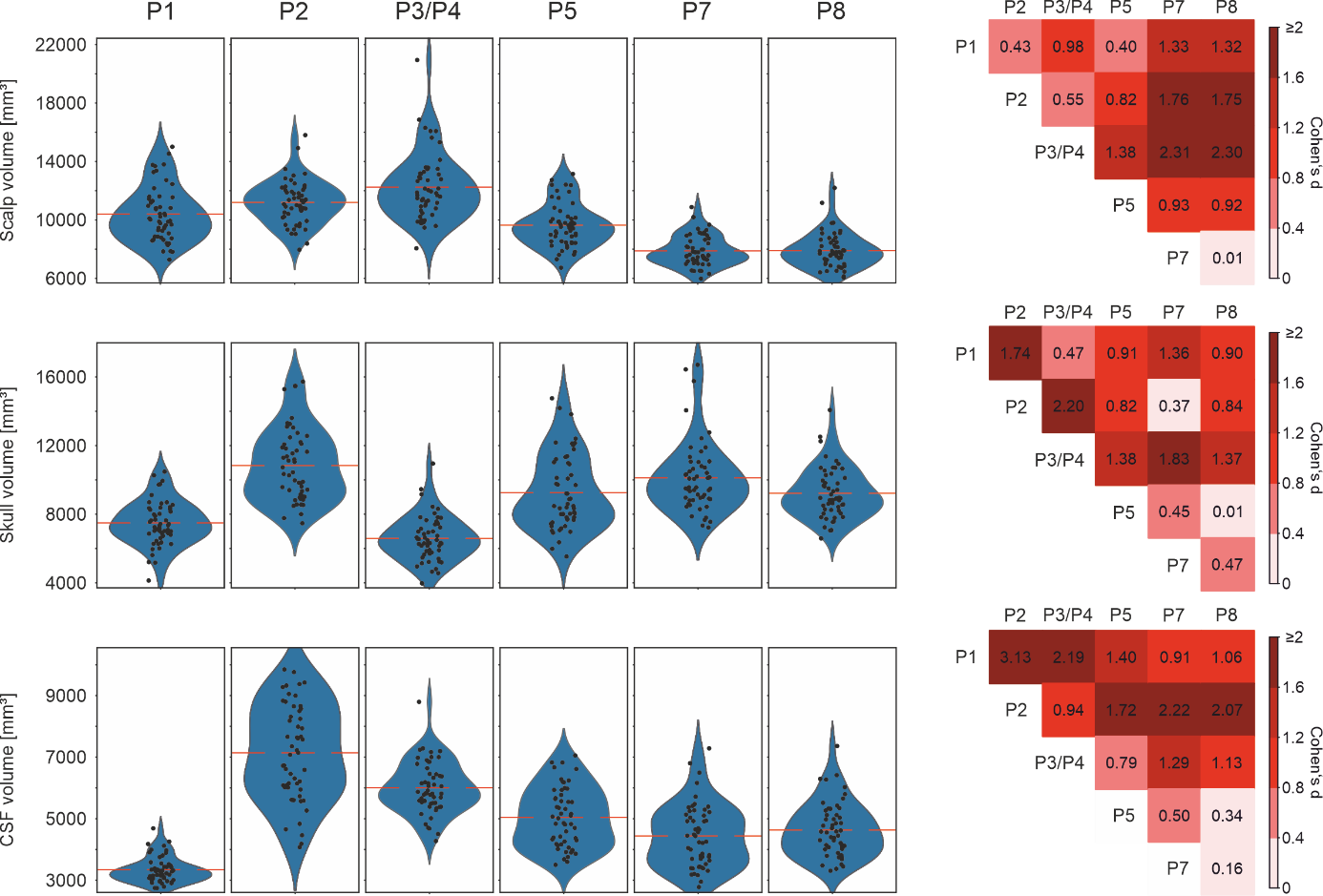

***Supplementary Figure 5.*** ROI sizes of scalp (A), CSF (C), and bone (E) area over the grey matter target regions and their pairwise comparison (B, D, F).

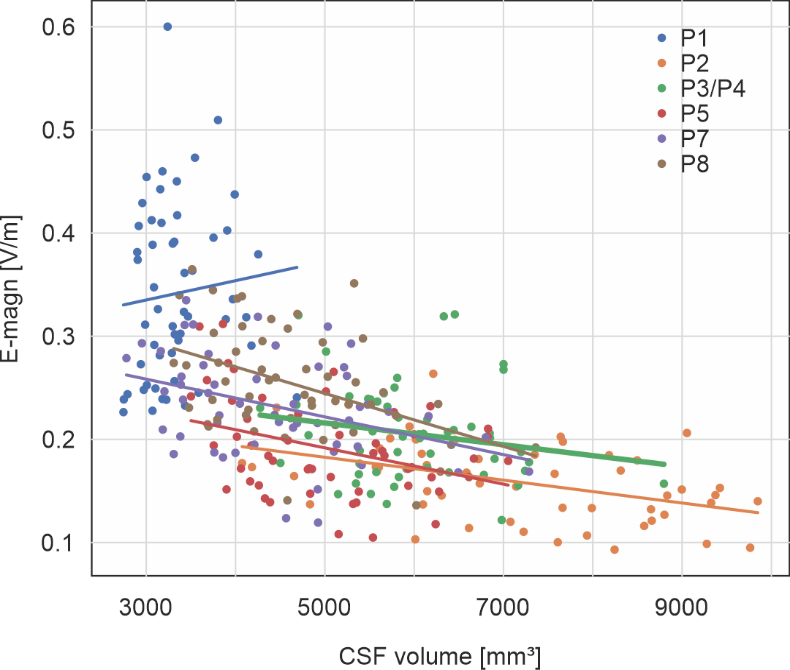

***Supplementary Figure 6.*** Scatterplot of electric field magnitudes and CSF volume for the cortical targets.

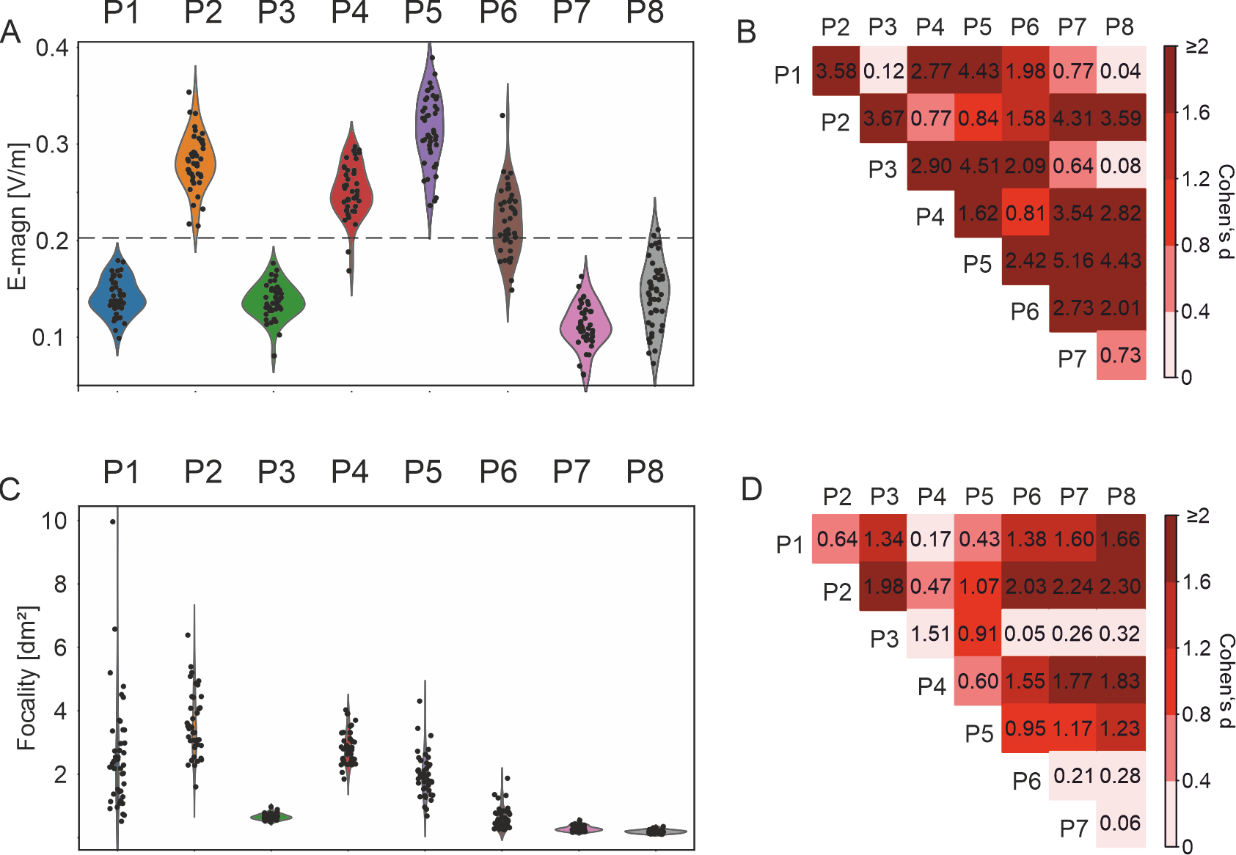

***Supplementary Figure 7.*** Conventional montages selected from the literature. Electric field magnitudes (A), and their pairwise comparison (B). Field focalities (C), and their pairwise comparison (D).

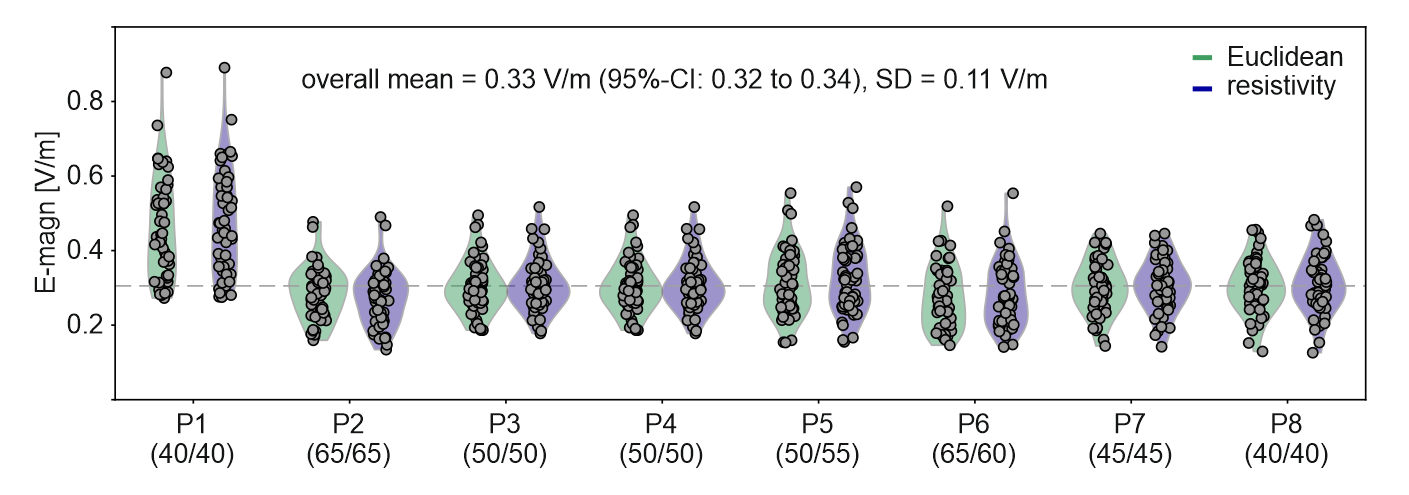
***Supplementary Figure 8*.** Electric field magnitude (95% percentiles) induced in target ROIs in 53 participants (Sample 2) with the resultant set-up for P1-8 (radii of Euclidean/resistivity algorithms in brackets). Even though the montages were optimized based on median of the field magnitudes rather than the 95% percentiles evaluated here, the 95% percentiles vary across the same overall mean in all projects except P1. This shows that our approach to harmonize the group-average field strength across ROIs performs robustly and is insensitive to the specific choice for evaluating the field magnitude in the ROIs. As expected from the results presented in the main paper for the median of the field magnitudes, also the 95% percentiles for P1 stayed higher than for the other projects. This is caused by our choice to maintain a montage radius of at least 40 mm to ensure practical feasibility.

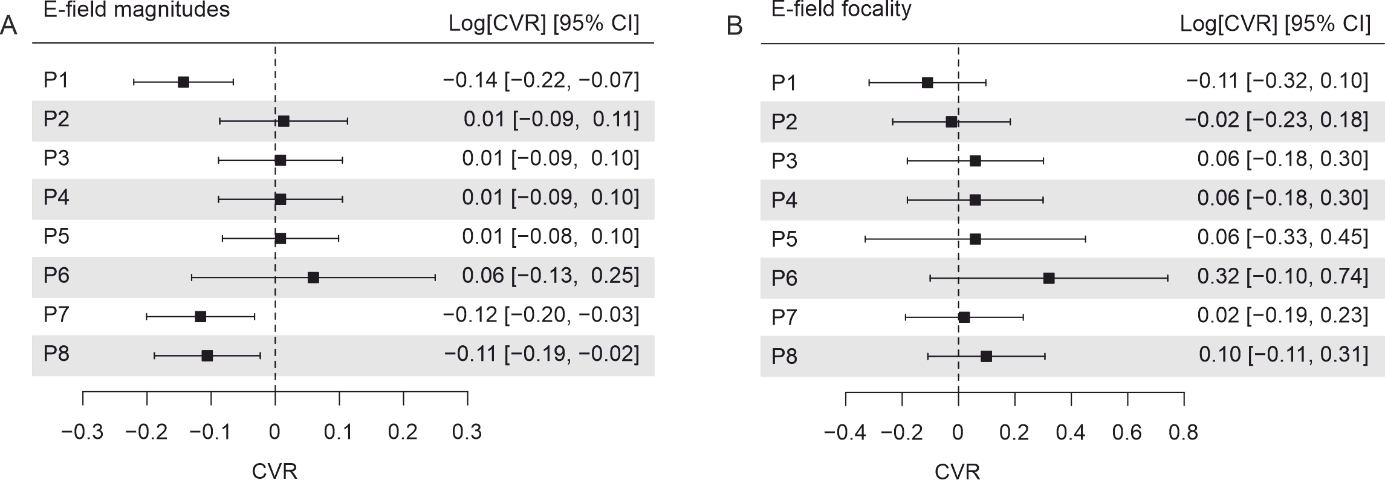

***Supplementary Figure 9.*** Comparison of variability between ROI of the projects for electric field magnitude (A) and focality (B). CVR, coefficient of variation.

**Supplementary Tables**

***Supplementary Table 1.*** Project and task details.

|  | Target cognitive/motor domain | Target brain region | Task administered during f-tES |
| --- | --- | --- | --- |
| P1 | Object-location learning | Right occipitotemporal cortex | associative learning of objects and their location on a schematic street map |
| P2 | Tacto-spatial working memory | Left posterior parietal cortex | retro cue delayed matching to sample task |
| P3 | Novel-word learning | Left inferior frontal gyrus | associative picture-pseudoword learning paradigm |
| P4 | Verbal working memory | Left inferior frontal gyrus | n-back task (remembering positions of letters in sequences with varying load) |
| P5 | Motor sequence learning | Left primary motor cortex | serial reaction time task |
| P6 | Eyeblink conditioning | Right cerebellum | delay eyeblink conditioning (using conditioning stimuli from different modalities, that is auditory and visual) |
| P7 | Learning-based control | Right dorsolateral prefrontal cortex | adapted face Stroop task (conflict-triggered adaptive control paradigm with name-face associations) |
| P8 | Value-based learning | Left dorsolateral prefrontal cortex | 3-state Markov decision task (two-alternative forced choice task between choice options in immediate and delayed conditions) |

***Supplementary Table 2.*** Parameters set in SimNIBS for the empirical montages.

|  | Current intensity | Anode | | | Cathode | | |
| --- | --- | --- | --- | --- | --- | --- | --- |
|  | *tdcs_list.currents* | *elect1.centre* | *shape* | *dimensions* | *elect2.centre* | *shape* | *dimensions* |
| P1 | 1e-3, -1e-3 | P8 | rect | 70, 50 | AF3 | rect | 70, 50 |
| P2 | 2e-3, -2e-3 | P3 | rect | 50, 50 | Right cheek | rect | 50, 50 |
| P3 | 1.5e-3, -.5e-3, -.5e-3, -.5e-3 | F7 | ellipse | 20, 20 | Fp1, FC5, F3 | ellipse | 20, 20 |
| P4 | 2e-3, -2e-3 | FC5 | rect | 50, 70 | AF4 | rect | 70, 50 |
| P5 | 2e-3, -2e-3 | C3 | rect | 50, 50 | AF4 | rect | 50, 50 |
| P6 | 2e-3, -2e-3 | I2 | rect | 50, 50 | Right buccinator | rect | 50, 50 |
| P7 | 1e-3, -1e-3 | F4 | ellipse | 25, 25 | F4 | ellipse | 115, 115 |
| P8 | 2e-3, -.5e-3, -.5e-3, -.5e-3, -.5e-3] | F3 | ellipse | 10, 10 | F1, AF3, F5, FC3 | ellipse | 10, 10 |
