## Supplementary Material B for "Harmonizing the stimulation dose of focal tDCS across target sites"

### Supplementary Material B - Positioning approach for focal tDCS: Performance evaluation

The positioning approach consists of two complementary steps: The center electrode is placed above the ROI center, using either the Euclidean or resistivity approach. In addition, the radius of the montage is reduced as much as possible while – on average in the group – it is ensured that the desired field strength is reached in the target. In the following, we evaluate the performance of these two steps.

We first test how well the two approaches for placing the center electrode ensure a good tradeoff between intensity in the target and focality. In line with the main paper, we define *target intensity* as the median electric field strength induced in the target ROI (in V/m). *Focality* is defined as the area on the gray matter central surface in which the target intensity is exceeded (in mm^2^). We evaluate the *intensity–focality trade-off* by calculating the ratio *target intensity/focality*. The ratio increases with increasing target intensity and higher focality (i.e., smaller cortical areas in which the field strength exceeds the target intensity). Conversely, lower target intensities or lower focality (i.e., larger cortical areas with fields above the target intensity) decrease the ratio*.*

Using Sample 1, we simulated the electric fields when systematically displacing the electrode montages along two orthogonal lines from their original positions above the target ROIs in steps of 7.5 mm (**Supplementary Fig. 10**). The optimal center-surround radii for each project were used. For each of the two lines, the displacement-vs-target intensity curves were centered and normalized, and subsequently averaged across individuals. In addition, cubic spline functions were fitted to the individual displacement-vs-target intensity curves to accurately estimate the displacement maximizing the target intensity using the spline function in Matlab. Focality and the intensity/focality ratio were analyzed in the same way.

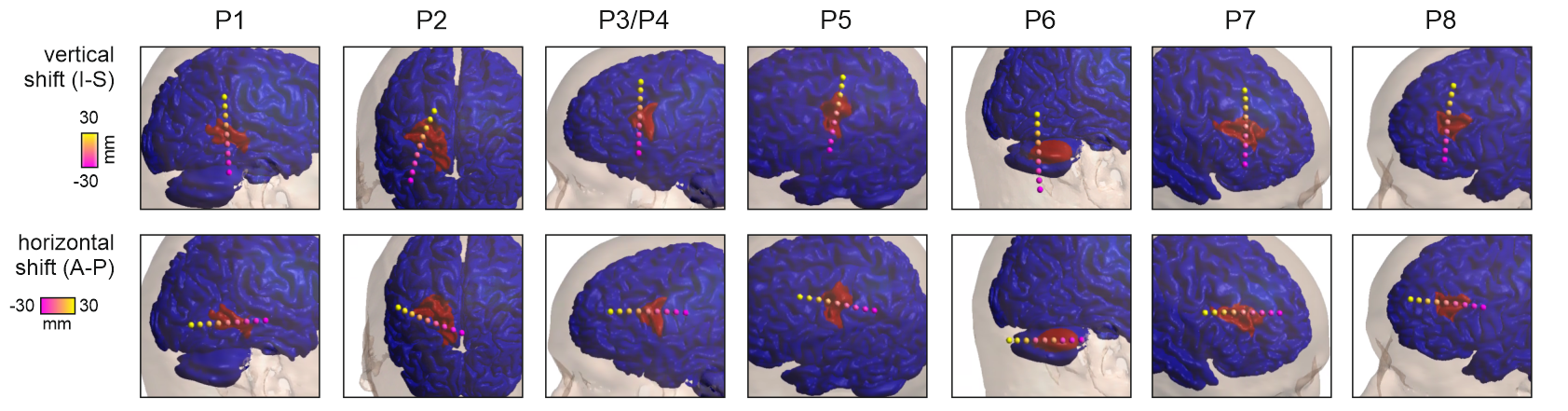

***Supplementary Figure 10.*** Illustration of the two orthogonal lines with shifted center electrode positions for an example head model. I-S, inferior-superior. A-P, anterior-posterior.

**Supplementary Fig. 11A** displays exemplar results for P3/P4, demonstrating that placing the center electrode above the ROI (displacement of 0) ensures a good intensity/focality tradeoff. This is particularly apparent for the A-P line, where the optimal position for the intensity/focality tradeoff stays close to 0 despite shifted curves for target intensity and focality. The results for the other target ROIs (see **Supplementary Table 3** for mean and SD of the optimal displacements across head models) generally confirm this conclusion. As exception, the optimal deviation for the cerebellum ROI in P6 reaches -12 mm along the A-P direction (**Supplementary Fig. 11B**). However, the curves for target intensity, focality and focality-intensity ratio remain rather flat for negative displacements between 0 and -15 mm in this case, so that the non-optimal placement at 0 is of limited practical relevance.

Second, we tested whether off-target stimulation is reliably minimized by reducing the radius of the montage as much as possible while keeping – on average in the group – the desired field strength in the target. For that, we plotted the dependence of focality on the montage radius for all projects (see **Supplementary Figure 12A**). As our focality measure scales with both the target intensity and the spread of the induced currents, we additionally calculated the gray matter area outside the target ROI in which the field strength exceeds a fixed threshold of 0.2 V/m (**Supplementary Figure 12B**). This confirms that the off-target stimulation decreases monotonically with decreasing montage radius.

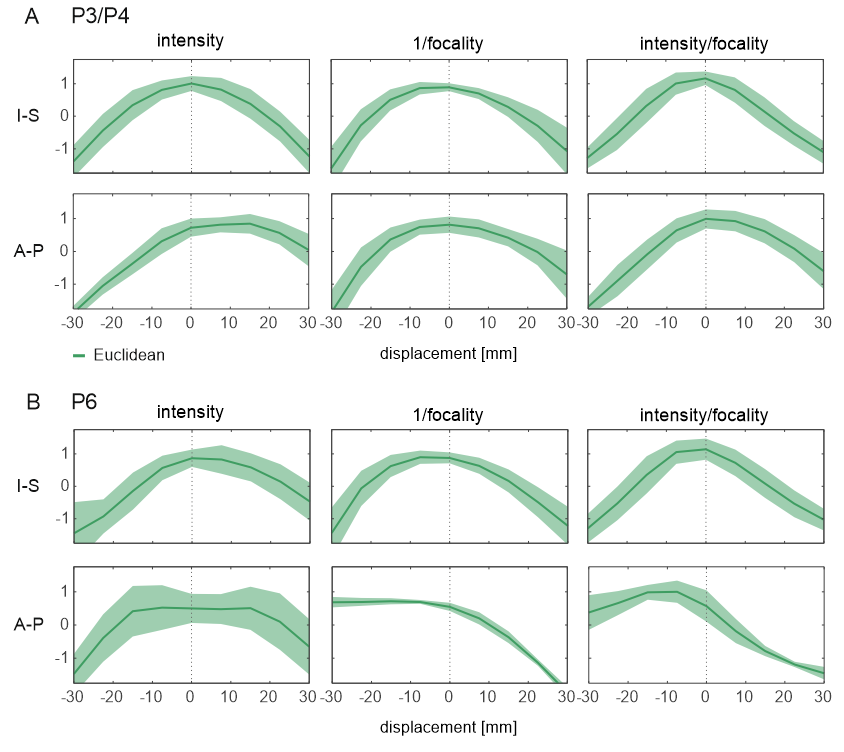

***Supplementary Figure 11.*** Results for P3/P4 and P6 (Euclidean approach) when displacing the montages along two orthogonal lines. (B) Group results for P3/P4 for target intensity (left), 1/focality (middle), and the ratio between target intensity and focality (right). The red lines show the group mean and green areas the SD. (C) Group results for P6, in which the optimal displacement for the A-P direction (-12 mm) deviated most strongly from 0. However, all three curves generally remain rather flat in the range from -15 mm to 0 mm, indicating that a non-optimal placement at 0 is of limited practical relevance.

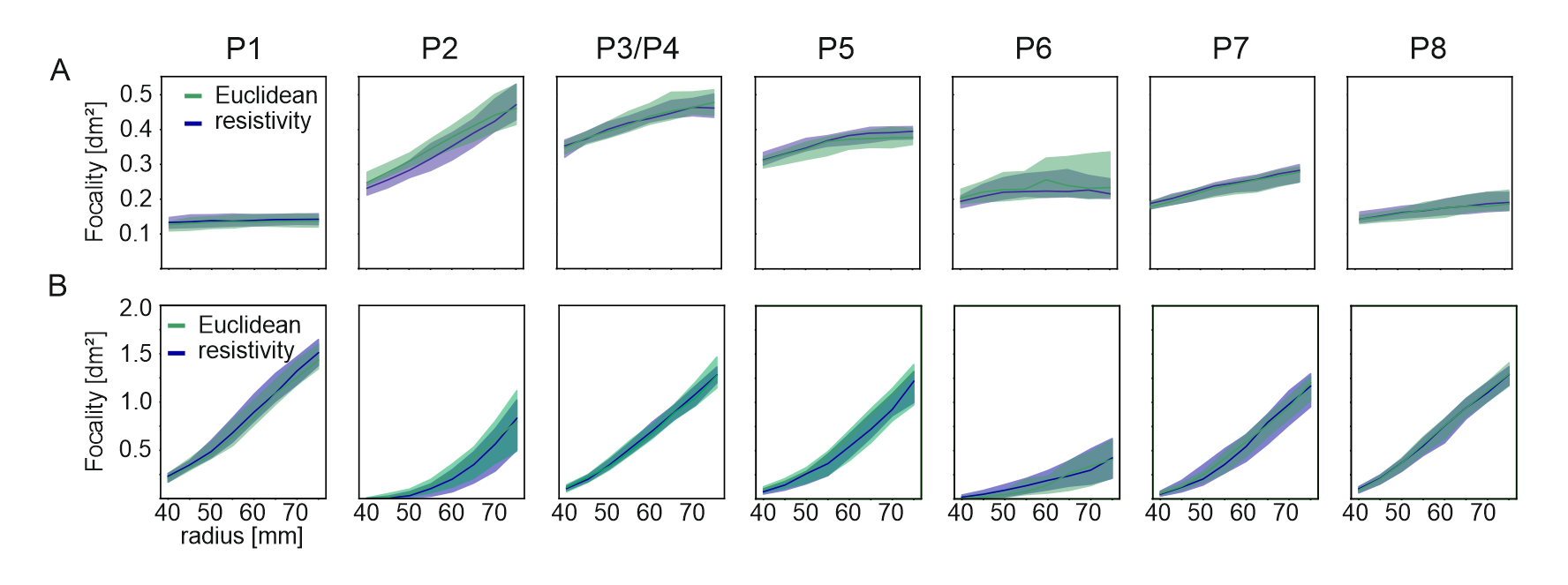

***Supplementary Figure 12.*** Focality in dependence of montage radius for the different target ROIs. (A) Focality is evaluated as the area of the gray matter central surfaces in which the field strength exceeds the median electric field in the target ROI. (B) Focality is calculated as the gray matter area outside the target ROI in which the field strength exceeds 0.2 V/m.

***Supplementary Table 3.*** Optimal displacements for all projects and the Euclidean or Resistivity approaches

|  |  | | | **Intensity** | | | | | | |  | | |
| --- | --- | --- | --- | --- | --- | --- | --- | --- | --- | --- | --- | --- | --- |
|  | Euclidean | | | | | |  | Resistivity | | | | | |
| P | Vertical shift (I-S),  Mean ± SD [mm] | | | Horizontal shift (A-P),  Mean ± SD [mm] | | |  | Vertical shift (I-S),  Mean ± SD [mm] | | | Horizontal shift (A-P),  Mean ± SD [mm] | | |
| 1 | -3.0 | ± | 6.5 | -0.5 | ± | 2.9 |  | -2.6 | ± | 4.7 | -1.1 | ± | 3.5 |
| 2 | 17.4 | ± | 7.8 | -14.3 | ± | 6.8 |  | 21.4 | ± | 6.7 | -16.4 | ± | 7.0 |
| 3/4 | -0.14 | ± | 7.2 | 8.8 | ± | 8.2 |  | -2.2 | ± | 6.7 | 7.4 | ± | 8.4 |
| 5 | 0.8 | ± | 8.0 | -12.9 | ± | 6.9 |  | 1.0 | ± | 8.6 | -17.6 | ± | 6.6 |
| 6 | 3.4 | ± | 10.1 | 2.8 | ± | 14.2 |  | 2.4 | ± | 7.0 | 4.0 | ± | 12.5 |
| 7 | -0.3 | ± | 7.5 | -4.2 | ± | 5.4 |  | -0.4 | ± | 7.2 | -4.6 | ± | 6.4 |
| 8 | 0.8 | ± | 7.2 | -0.6 | ± | 5.9 |  | 0.8 | ± | 7.7 | 0.4 | ± | 5.6 |
|  |  |  |  | **Focality** | | | | | | |  |  |  |
| 1 | 3.0 | ± | 6.1 | -4.0 | ± | 4.9 |  | 6.1 | ± | 6.4 | -5.2 | ± | 6.0 |
| 2 | -2.8 | ± | 6.1 | 10.4 | ± | 6.2 |  | 0.3 | ± | 5.2 | 6.6 | ± | 6.3 |
| 3/4 | -2.2 | ± | 5.3 | -0.5 | ± | 8.4 |  | -4.0 | ± | 5.3 | -1.5 | ± | 9.2 |
| 5 | 0.3 | ± | 5.7 | 8.4 | ± | 9.6 |  | -1.6 | ± | 7.0 | 4.3 | ± | 11.7 |
| 6 | -4.2 | ± | 6.5 | -15.6 | ± | 10.6 |  | -6.4 | ± | 6.6 | -15.7 | ± | 11.8 |
| 7 | -1.8 | ± | 5.4 | 3.5 | ± | 4.4 |  | -2.0 | ± | 6.2 | 2.8 | ± | 5.2 |
| 8 | 2.3 | ± | 3.2 | 2.8 | ± | 4.7 |  | 1.5 | ± | 3.9 | 3.5 | ± | 5.1 |
|  |  |  |  | **Intensity/focality trade-off** | | | | | | |  |  |  |
| 1 | 1.3 | ± | 3.9 | -2.3 | ± | 3.1 |  | 2.9 | ± | 4.6 | -3.1 | ± | 4.1 |
| 2 | 1.7 | ± | 6.7 | 6.7 | ± | 5.8 |  | 4.9 | ± | 5.2 | 2.0 | ± | 6.4 |
| 3/4 | -1.8 | ± | 5.4 | 2.7 | ± | 7.2 |  | -3.8 | ± | 4.8 | 1.9 | ± | 8.2 |
| 5 | 0.6 | ± | 5.4 | 1.3 | ± | 6.5 |  | -0.5 | ± | 6.8 | -3.3 | ± | 7.5 |
| 6 | -2.7 | ± | 5.7 | -12.3 | ± | 8.7 |  | -2.9 | ± | 5.6 | -10.2 | ± | 8.6 |
| 7 | -2.0 | ± | 4.7 | 1.5 | ± | 4.0 |  | -1.6 | ± | 5.4 | 1.0 | ± | 4.9 |
| 8 | 2.4 | ± | 2.8 | 2.0 | ± | 3.7 |  | 1.8 | ± | 3.5 | 2.8 | ± | 4.6 |
